# Changing abiotic and biotic environments affect the genetics of adaptation and speciation

**DOI:** 10.64898/2026.07.15.738782

**Authors:** Trey J. Scott, David C. Queller

## Abstract

Abiotic and biotic environments can change in different ways that affect how populations adapt, diverge, and speciate. However, many models of speciation only examine adaptation to constant environments. We used Fisher’s geometric model to study adaptation and speciation in different types of changing abiotic and biotic environments. We hypothesized that, when the magnitude of environmental change is equal, fitness would be lowered more by consistently directional abiotic change and by consistently antagonistic biotic conflict. As a result, directional changes and biotic conflict would cause a greater evolutionary response through fixation of adaptive mutations, greater divergence between populations, and lower migrant or hybrid fitnesses (higher speciation potential). Increasing the directionality of abiotic change did indeed have all these effects. However, populations evolving with conflict, despite showing the expected arms-race elevation of deleterious change and evolutionary response, did not result in decreased fitness of migrants and hybrids. Two factors seem to be involved. In tug-of-war conflict, high total phenotypic change does not lead to high phenotypic divergence because the phenotypic change churns around in the same general area of trait space. With victim-exploiter (trait matching) conflict, hybrids and migrants were frequently more fit than those with other kinds of change because of escape from locally adapted antagonists. In this case, foreign genes sometimes allowed populations in conflict to generate useful variation. These results show that different kinds of ecological change can affect adaptation and the formation of new species in distinct ways and that local divergence and incompatibility do not always go hand in hand.

**Significance Statement:** Adaptation and speciation are key processes in evolution. During the process of adapting, populations genetically diverge from each other and can become incompatible, leading to new species. The tendency for populations to diverge can be different in changing environments. Changes in the environment can consist of changes in the non-living parts like temperature or elevation (the abiotic environment) or changes in the species that a population interacts with (the biotic environment). Abiotic and biotic environments are thought to change in different ways because biotic environments are evolving but abiotic ones are not. In particular, antagonistic biotic evolution can lead to arms races. We modeled how abiotic change and biotic changes from antagonistic or mutualistic interactions affect adaptation and speciation. We found that, as expected, antagonistic biotic changes led to a large amount of evolutionary divergence but surprisingly this did not lead to correspondingly large reduction in migrant or hybrid fitness. In a tug-of- war antagonistic model, this was due evolutionary churn – high change did not lead to high divergence. In a victim-exploiter (trait matching) model, migrants and hybrids between diverging populations were often better off because antagonisms were broken up. As a result of migrants and hybrids having higher fitness, antagonistic biotic interactions may result in less speciation than expected from their evolutionary arm-races.

## Introduction

Different kinds of biotic and abiotic environmental changes, or “environmental change regimes”, are expected to affect how a population adapts [1–7]. These differences in adaptation will be reflected in the kinds of mutations that are fixed, the genetic architecture of evolving traits, and fitness. We call such changes a population’s “adaptive response.” For example, in a meta-analysis of plant quantitative trait loci, adaptation to biotic environments involved fixing mutations with larger phenotypic effects compared to adaptation to abiotic environments [2]. Though some, mostly theoretical, studies have found different adaptive responses between some kinds of changing environments [2,3,8,9], the general rules of adapting to changing environments are not fully clear [4,5]. Still less clear is how differences in adaptive responses may then affect divergence between populations and lead to speciation except for an expectation that more adaptive change should lead to more speciation [10,11].

Adaptive responses may be the most different between populations adapting to biotic and abiotic regimes. Biotic interactions can involve either partners that are mutualistic or partners that are in conflict, such as predator-prey or pathogen-host interactions. Conflict is thought to result in strong and persistent maladaptation due to arms races, where each party has to respond to the antagonistic adaptations of the other [12], while having consistently low fitness over time [3]. Low fitness allows for an increased capacity to adapt through the fixation of large and frequent substitutions [3], which might be expected to lead to increased divergence and speciation.

Arms races can be conceptualized with a joint phenotype. A joint phenotype is a single trait that is shared by two or more parties that jointly affect it [6,13]. For example, the amount of leaf eaten by an herbivore is a joint phenotype since the plant can affect this phenotype by evolving toxins or other defenses while the herbivore can evolve resistance. Tug-of-war arms races can result if the optimum values for the amount of leaf eaten differs between the plant and the herbivore.

Antagonistic partners will never agree on the optimal value of the joint phenotype, so they will evolve adaptations and counter-adaptations in an arms race [3,6]. This means that biotic conflict will tend to result in environmental changes that are consistently detrimental and keep both parties in a state of rapid change and maladaptation called a Sisyphean arms race [3].

In contrast to arms races, abiotic changes to the optimum are expected to be more random with respect to the fitness interests of the adapting population. Other things being equal, this randomness is expected to result in a smaller amount of evolutionary change and maladaptation [3].

A simple theoretical model has shown that these differences in maladaptation between biotic and abiotic change can occur and have important impacts on a population’s adaptive response [3], but more detailed modeling is needed to fully understand the differences in adaptive responses between abiotic and biotic regimes and how abiotic and biotic changes affect speciation.

The earlier model comparing abiotic to biotic change focused on random abiotic changes [3]. However, abiotic changes in some environments may be more likely to resemble past changes and exhibit temporal correlations [14]. These correlations could result in abiotic environmental change that moves in a consistently deleterious direction, like the effects of biotic conflict.

On the biotic side, the previous model assumed that biotic interactions are harmful and investigated only tug-of-war evolution [3]. Joint phenotype conflict can also be modeled with explicit trait interactions that can involve a chase rather than a pull. For example, some interactions involve matching between traits, where an exploiter species benefits by matching or unlocking the traits of a victim species [15,16]. This victim-exploiter chase conflict could result in different arms race dynamics than the Sisyphean arms races from a tug of war.

Beyond conflict, biotic changes can be beneficial, as in mutualisms. Mutualistic adaptation might lead to partners co-adapting and less strong adaptive responses. Using this more complete set of biotic and abiotic models could provide new insights into how adaptive responses differ between these environmental change regimes.

Finally, previous modelling has not fully explored the effects of different environmental change regimes on reproductive incompatibility and speciation. Speciation occurs when populations diverge so that members of different populations can no longer exchange genes [10]. Genetic exchange will be reduced when a population fixes alleles that are adaptive for its local environment, including its local gene pools, but maladaptive in the environment or gene pool of another population [11,17].

Both biotic and abiotic environmental changes can cause genetic divergence and ecological speciation [18]. An example involving largely abiotic selection comes from populations of insects that have adapted to different thermal niches and speciated along altitudinal gradients in tropical mountains [19]. Speciation can also be impacted by biotic environmental changes, especially those involving antagonistic partners [20–24]. For example, biotic interactions between butterflies and plants have long been thought to be important drivers of diversification [25] though empirical evidence is mixed [26–28]. Biotic interactions may also increase the number of diverging loci that lead to incompatibilities because of arms race dynamics [21,29]. Even beneficial biotic interactions can be important for speciation, as in the case of partner switching in plants and their pollinators [30,31].

One key assumption of most speciation models is that diverging populations become well adapted to their local environments and maladapted in foreign environments. However, maladaptation appears to be common even in home environments [32] and can result from changing biotic and abiotic environments [3]. Maladaptation to the home environment may diminish the relative disadvantage for migrants and hybrids.

The amount and nature of divergence between nascent species may be affected by the genetics of adaptation. The number and size of fixations that differ between populations may be important [33] and is thought to differ when populations experience different kinds of changing environments [3]. Effects of individual genes are often thought to be small but need not be. For example, a single allele in monkeyflower can have a large effect on whether a plant is visited and pollinated by bees or hummingbirds [34]. Whether a gene has deleterious pleiotropic effects can also affect speciation [35,36].

Fisher’s [37] geometric model has emerged as a powerful model for understanding adaptation to different changing environments [3,8,38–41] and the genetics of speciation [35,42–47]. The standard geometric model without environmental change, where a population adapts to a constant environment, assumes a monomorphic population with *n* quantitative traits that are affected by stabilizing selection centered at some optimal phenotype. A population adapts to this optimum by successively fixing mutations that move the population from a lower fitness phenotype to a higher fitness phenotype closer to the optimum, though this can be opposed by drift [48]. Because of the geometry of this process, mutations are more likely to fix and fixations will have larger effects when populations are farther from their optima — when they are more maladapted.

Another consequence is that large mutations will fix during the initial stages of adaptation, but in later stages will more often overshoot the optimum and be disfavored [48]. Though simple, the geometric model has been used to address myriad questions in population genetics [49] and has proven consistent with many empirical patterns [43,44,50].

The geometric model can recapitulate major findings of hybridization studies like the decrease in F2 hybrid fitness with increasing genetic distance between parents (F2 speciation clock) and heterosis [43]. Hybridization is achieved in the model by taking two parental populations, which have independently fixed alleles in their local environments (these environments can have the same or different optima), and combining a random sample of their fixations to construct hybrid offspring phenotypes [42]. Fitness of hybrid crosses can be measured in both parental environments to understand the role of ecology in leading to ecological speciation [35]. Another way that speciation occurs that can be modeled with the geometric model is when migrants have lower fitness in new environments. This has the same consequences as low fitness hybrids on speciation except that it prevents hybrids from being produced in the first place [51].

The tendency to speciate should be affected by differences between conflict, abiotic change, mutualisms, and adaptation to a constant optimum. Isolated populations will fix numerous mutations and become genetically diverged. Many lineage-specific fixations may be incompatible in other lineages and contribute to speciation. Thus, as a first hypothesis, 1) environmental regimes that are more detrimental to fitness should cause adapting populations to evolve more, 2) greater evolution should cause adapting populations to become more divergent, and 3) greater divergence should cause adapting populations to produce low fitness hybrids and migrants (Fig 1).

**Fig 1:**
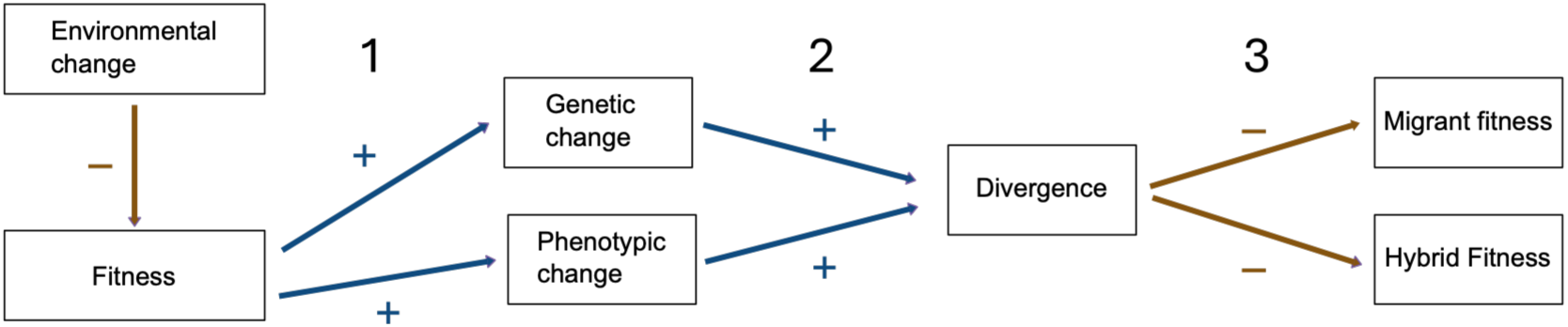
Schematic of the causal chain for how environmental effects on fitness should lead to incipient speciation. Plus and minus signs show direction of hypothesized correlation. For example, more environmental change should lower fitness and cause 1) more genetic change, 2) more phenotypic divergence, and 3) lower migrant and hybrid fitness.

Because of the tendency for conflict to result in arms races, it has often been suggested that populations experiencing conflict should have even higher genetic divergence and more incompatibilities [6,21,29,52,53]. Increased divergence is a common outcome of coevolution [54–57], but evidence that this leads to more incompatibilities is more sparse [22,58]. For mutualisms, stabilizing selection may dominate and lead to low divergence and speciation [59]. For abiotic change, we might expect greater incompatibility in highly directional regimes where environmental changes are correlated in direction, for example consistently getting hotter.

We use the geometric model to investigate how different kinds of environmental change (environmental change regimes) affect a population’s adaptive response and how these adaptive responses lead to speciation. We first quantify how different kinds of environmental change lead to different adaptive responses. We then test migrants and hybrid crosses between two independently evolving populations experiencing the same type of environmental change to determine how different kinds of environmental change affect the tendency to lower migrant and hybrid fitness.

## Overview of simulations

To understand speciation due to different adaptive responses to kinds of environmental change, we developed an R package called *SimGM* (https://gitlab.com/treyjscott/SimGM) to simulate adaptation to different changing environment regimes (the basic models are shown in Fig 2 and described in Table 1; see methods for more details). Briefly, these regimes include standard adaptation to a constant optimum, adaptation to a changing abiotic environment (with varying temporal correlations measured by the correlation coefficient *r*), coevolution with antagonistic partners (modeled with both tug-of-war and victim-exploiter conflict), and coevolution with mutualistic partners. In our simulations of tug-of-war conflict and mutualisms, both partners evolve such that partners are interchangeable (both partners experience symmetric selection) so we arbitrarily select one partner to use in our results. However, victims and exploiters evolve in distinct ways (selection is asymmetric) so we track both victims and exploiters in our results. Thus, there are four biotic change regimes: tug of war, victim, exploiter, and mutualism.

**Fig 2:**
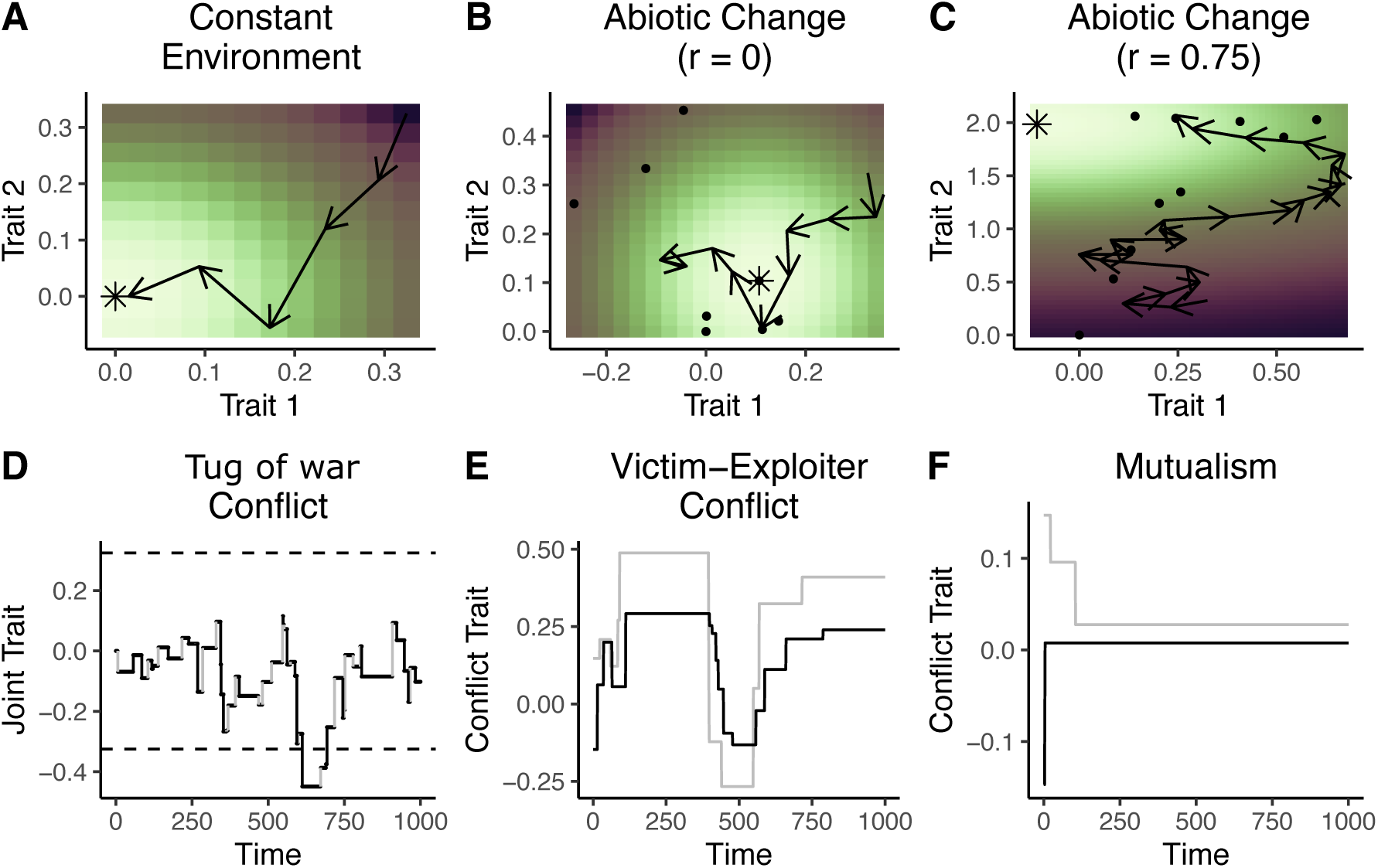
Example simulations of models of trait change under different models of environmental change. The first row shows changes in two dimensions (A) constant environment, (B) uncorrelated abiotic change (*r* = 0), and (C) highly correlated abiotic change (*r* = 0.75). Colors show fitness (for changing environments colors reflect fitness based on the final optimum position) with purple being low fitness and yellow being high fitness. Arrows show the effect of mutations changing the ancestral phenotype to the mutant phenotype. The second row shows biotic simulations (shown with one trait for simplicity). From left to right, plots show (D) tug-of-war conflict over a joint trait where both parties can shift the value of the trait, but have different optimal values (dashed lines), (E) victim-exploiter conflict where exploiters (black) chase victims (gray), and (F) mutualism where both benefit by matching their traits.

**Table 1:**
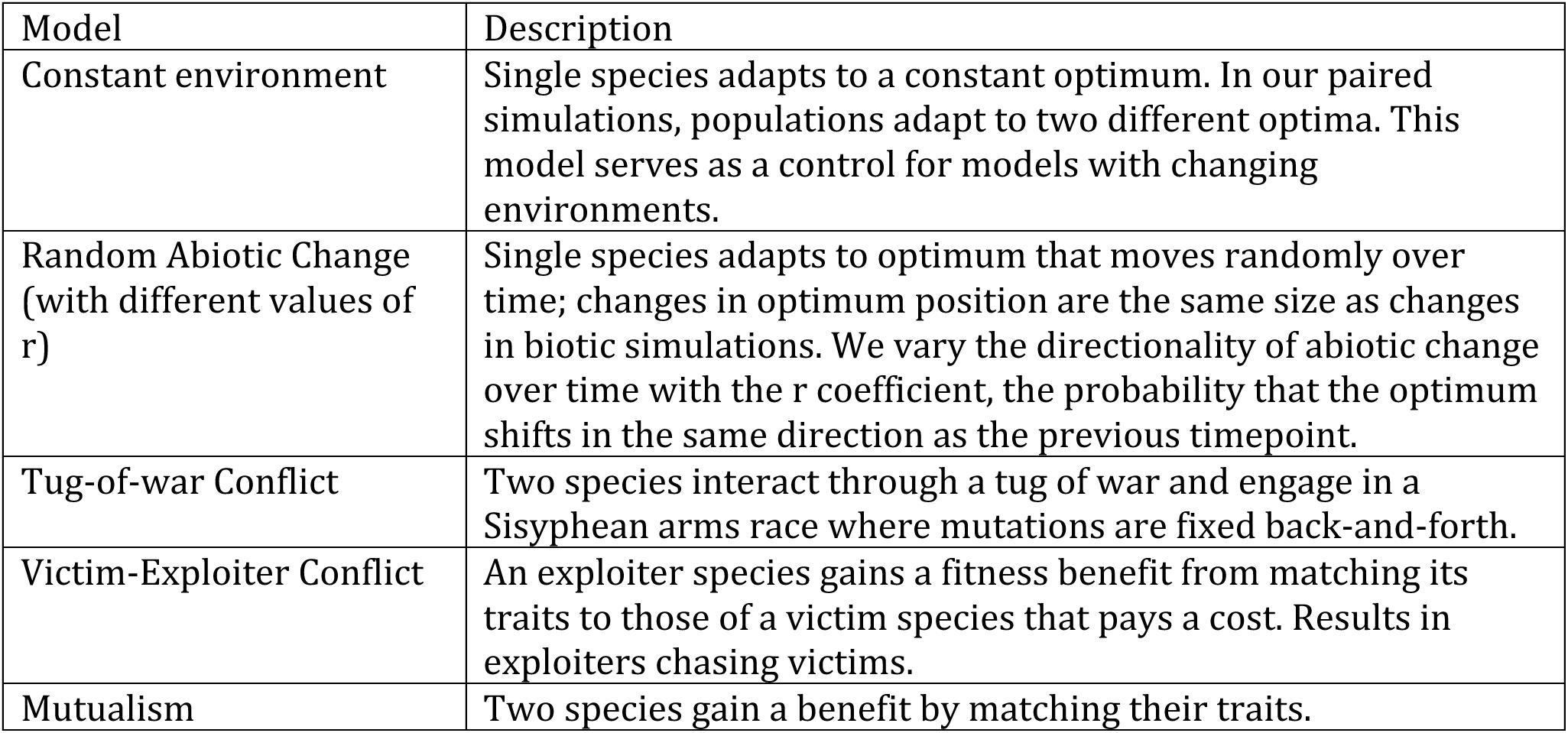
Environmental change regimes.

| Model | Description |
| --- | --- |
| Constant environment | Single species adapts to a constant optimum. In our paired simulations, populations adapt to two different optima. This model serves as a control for models with changing environments. |
| Random Abiotic Change (with different values of r) | Single species adapts to optimum that moves randomly over time; changes in optimum position are the same size as changes in biotic simulations. We vary the directionality of abiotic change over time with the r coefficient, the probability that the optimum shifts in the same direction as the previous timepoint. |
| Tug-of-war Conflict | Two species interact through a tug of war and engage in a Sisyphean arms race where mutations are fixed back-and-forth. |
| Victim-Exploiter Conflict | An exploiter species gains a fitness benefit from matching its traits to those of a victim species that pays a cost. Results in exploiters chasing victims. |
| Mutualism | Two species gain a benefit by matching their traits. |

In nature, different kinds of environmental change may occur on different timescales and involve different magnitudes of change. However, we wanted to abstract away from these differences to compare biotic and abiotic regimes that experience the same degree of environmental change. We can then identify differences due primarily to the sign of environmental change, which is typically negative in antagonistic interactions, positive in mutualistic ones, and variable in abiotic interactions (increasingly negative with correlated or directional changes). To do this for each biotic simulation, we performed a parallel abiotic simulation with the same timing and magnitude of change, but not necessarily the same direction of change (see Methods and [3]). We did this for each kind of biotic change (tug of war, victim, exploiter, and mutualism) with each level of abiotic temporal change correlation (*r* = 0, 0.25, 0.5, 0.75, 1). Increasing *r* results in more directional change (see Methods and Fig 2). This means that for each biotic change regime, there were five abiotic change simulations with identical timing and magnitudes of change that differed in directionality of change.

We label our abiotic change simulations according to the first letter of the source of their corresponding biotic change and the amount of correlation. For example, “abiotic M0.5” is abiotic change where the magnitude of changes corresponds to that which a mutualist experiences and has a temporal correlation of 0.5. Altogether, our models of changing environments result in a set of 25 distinct environmental change regimes that we tested (standard adaptation, four kinds of biotic change, and 20 kinds of abiotic change). Importantly, because we force the frequency and magnitude of biotic and abiotic change to be equal, differences in our models between abiotic a biotic speciation would indicate some important difference other than magnitude and frequency of change.

Adaptation in the geometric model can be affected by different parameterizations like the initial fitness of a population, the strength of stabilizing selection, and the number of traits [3,49]. To account for the effect of different model parameterizations, we ran replicate simulations with randomly selected parameters from uniform distributions within ranges given in Table 2. Within each replicate, each of our 25 environmental change regimes had the same parameter values. This created blocks with the same parameterization, with the only difference being the environmental change regime.

**Table 2:** Parameters that were varied in simulations used to generate adaptive responses and construct hybrid crosses.

| Parameter name | Range | Description |
| --- | --- | --- |
| $a$ | 0.1-1.5 | Strength of stabilizing selection |
| $\bar{m}$ | 0.01-0.2 | Average size of mutations |
| $L$ | 0.075-0.2 | Initial lag load of populations that determines the potential for adaptation at the start of simulations (also scales the amount of initial conflict) |
| $n$ | 1-4 | Total number of traits. We limited the number of traits because of the tendency for victims to escape exploiters with high dimensionality as demonstrated in [60]. |
| $v$ | 1-4 | Number of traits that are responsive to selection from environmental changes. $v$ determines the number of unresponsive traits ($p = n-v$). Unresponsive or pleiotropic traits do not respond directly to environmental changes but can respond via pleiotropy with the selected traits. |
| *Time* | 1500-15000<br>(increments of 500) | Number of simulated time points |

To measure adaptive responses to each environmental change regime, we calculated several summary measures of the outcome of adaptation from completed simulations. These summary measures, to be discussed later, reflect quantities that can be measured empirically.

To investigate the process of speciation, we generated additional simulations with identical parameters. This involved paired simulations, representing two newly isolated populations, that started with the same initial values and then diverged both genetically and environmentally (except for standard adaptation, where environmental divergence is set at the start of simulations so paired populations adapt to two different constant optima, but under the same environmental regime). We assessed migrant fitness by arbitrarily assigning one member of a pair to be the local population while the other member is the foreign population and assessed the fitness of a foreign genotype in the local environment. For hybrids, we cross parents from the local and foreign populations (see Methods) and assess hybrid fitness in the local parental environment.

## Results

### Directional abiotic change and biotic conflict cause exaggerated adaptive responses

Different environmental change regimes may engender different adaptive responses that could affect speciation. We will first discuss the differences in adaptive response for individual populations and then return to differences in speciation that occur between individuals from different populations. We focused on five components of a population’s adaptive response: the average fitness, the average fixation size, the fixation rate (proportion of mutations that fix), the average selection coefficient of fixed mutations (selection), and pleiotropic change (the average distance that traits that are unresponsive to changing environments are moved away from their fixed optima). See the Methods for more details. To determine if environmental change regimes affect adaptive responses, we performed principal components analysis and calculated the centroid position for each kind of environmental change.

We found distinct patterns in how our environmental change regimes clustered (Fig 3A). Points ranged largely along the diagonal, reflecting the fact that the first two principal components had similar loadings, with decreasing average fitness and increasing other components of the adaptive response, except pleiotropic change for PC2 (inset in Fig 3A shows loadings while Fig 3B-E shows individual components plotted against fitness). This inverse relationship – lower fitness being associated with higher adaptive response – is expected and has been shown in past models [3,49]. Mutualism, abiotic change based on mutualisms, and adaptation to a constant environment cluster on the lower left corner, with high fitness and low adaptive response, while simulations with biotic conflict and their respective abiotic change simulations are scattered to the right and upward, with lower fitness and higher adaptive responses (Fig 3A-E). As expected, within any type of abiotic change (T, V, E, M), higher directionality lowers fitness and increases adaptive response. Also as expected, simulations involving biotic conflict (tug of war, victim, exploiter) showed a similar lowering of fitness and increased adaptive response (Fig 3B-E).

**Fig 3:**
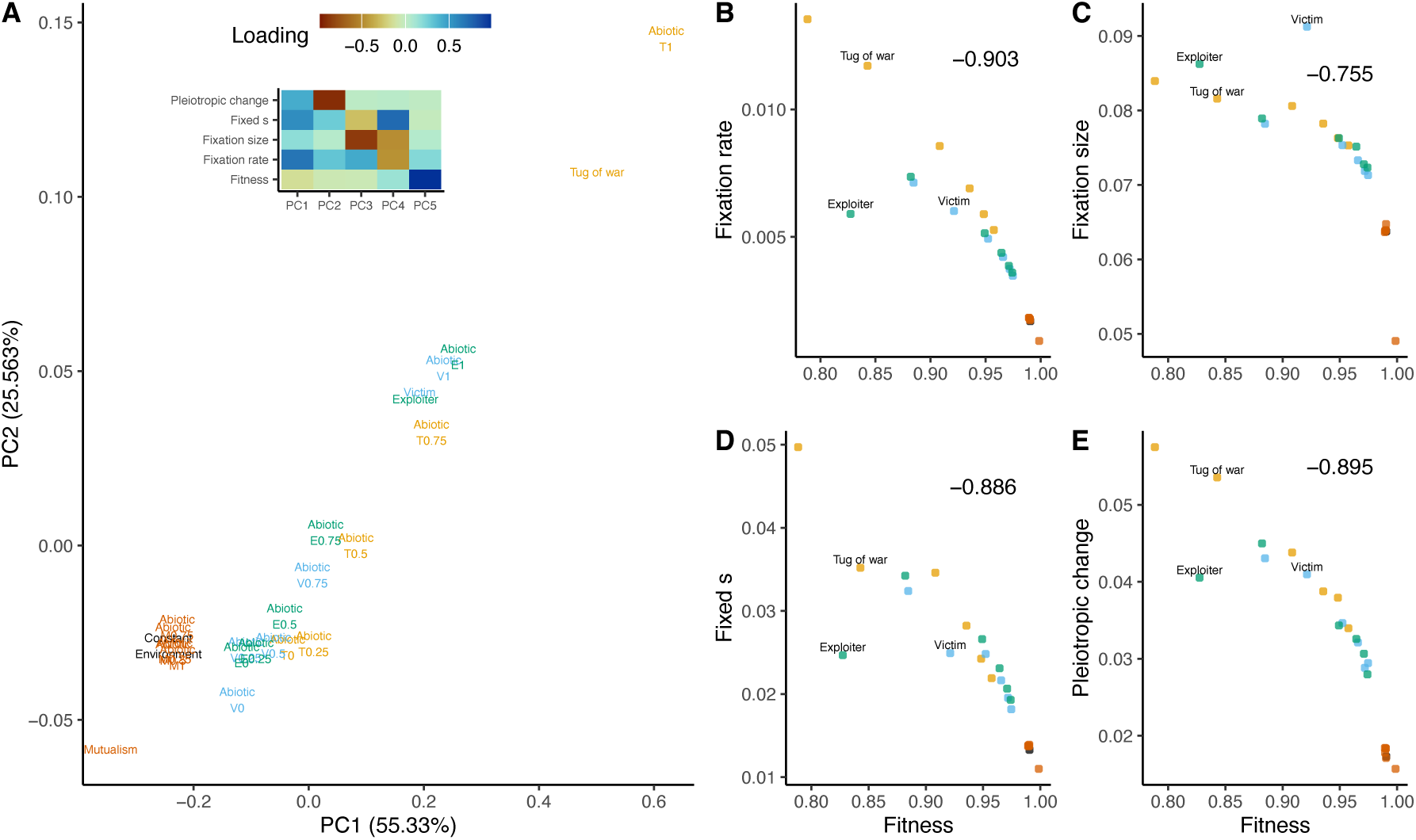
Different environmental change regimes have different adaptive responses. (A) centroids of different environmental change regimes in the space of the first two principal components. The names of different regimes show where the centroids are located. Colors distinguish biotics along with their respective abiotic comparison simulations that have all experienced the same magnitude of environmental change. The inset shows the loadings from the principal components analysis for the components of the adaptive response. (B-E) Average components of the adaptive response for all environmental change regimes plotted against the average fitness. Regimes experiencing conflict with exaggerated adaptive responses are labeled. Pearson correlations are shown in the top right corner for each plot in C-F.

### Directional change and biotic conflict are usually deleterious

A key way that abiotic change is thought to differ from conflict is that conflict results in environmental changes that are consistently harmful [3]. However, directional abiotic change may be harmful for the same reason. On the other hand, the pure mutualism in our models (with no conflict) should almost entirely experience beneficial environmental changes. We tested these predictions by calculating selection coefficients for individual environmental changes (due to shift in the optimum position for abiotic changes, or trait values of the partner for biotic changes). These differ from the selection coefficients in the last section, which were for fixed mutations, not environmental changes. We call the distribution of these effects during an adaptive walk the distribution of fitness effects of the environment (DFEE). This is analogous to the distribution of fitness effects (DFE) of new mutations that is commonly studied with the geometric model [41,61,62]. Just as the DFE measures fitness changes due to mutations, the DFEE measures the fitness change resulting from environmental changes. We ran simulations and calculated DFEEs for all environmental change regimes, except for those adapting to constant environment where there was no environmental change.

We found that environmental change, whether biotic or abiotic, tends to be deleterious, as measured both by the proportion of the DFEE that was negative (more than half of environmental changes lower fitness; Fig 4A) and by the median of the DFEE (Fig 4B). Under abiotic change, higher directionality consistently caused more negative effects on fitness. Biotic conflict generally caused more negative effects than abiotic change, with the exception of abiotic change that was highly directional. Mutualism, which we expected to be the opposite of conflict, showed almost no deleterious changes and largely positive median DFEE values while the corresponding mutualism abiotics were similar to other abiotics (Fig 4A&B).

**Fig 4:**
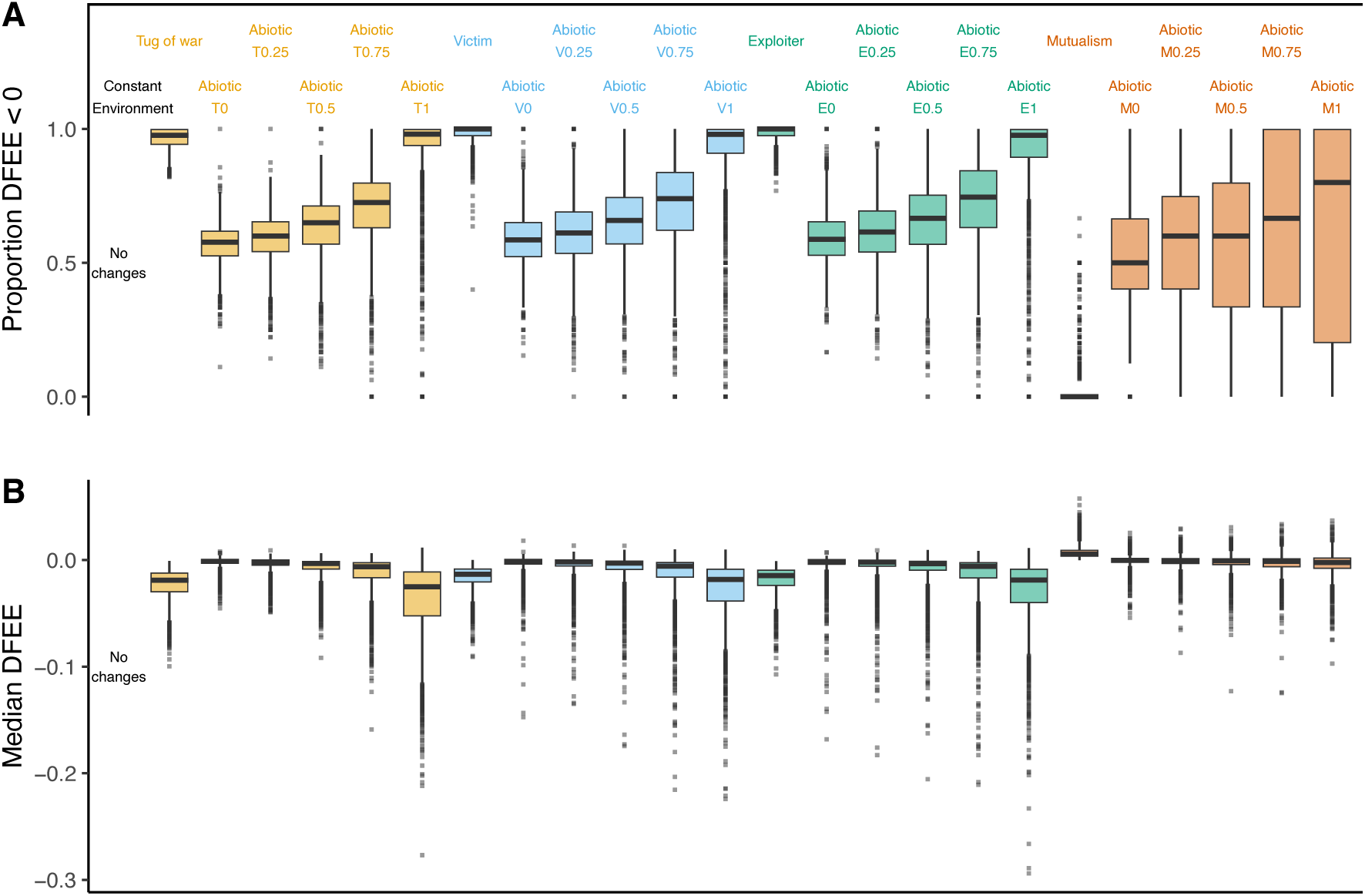
Environmental change involving conflict resembles directional abiotic change and is maladaptive. Plots show summary statistics of the distribution of fitness effects of the environment (DFEE) for tug-of-war, victim, exploiter, and mutualism regimes (along with corresponding abiotic change models) with varying directionality. A) The proportion of environmental changes that are deleterious for fitness (<0). B) Medians of the distribution of fitness effects of environmental change (DFEE). Colors distinguish biotics along with their respective abiotic comparison simulations that have all experienced the same magnitude of environmental change. Boxplots show the median line with the ends of the boxes corresponding to the first and third quartiles (25^th^ and 75^th^ percentiles, respectively). Lines show either the entire range of the data or, if there are outliers (shown as points), 1.5 times the range between the first and third quartiles.

Together, these DFEE results confirms that change due to abiotic conflict and highly directional abiotic change are more deleterious than mutualism and less directional abiotic change.

### Conflict and directional change cause more evolution and, except for tug of war, more divergence

Speciation occurs between populations that have diverged and become locally adapted to different environments. To understand how different environmental change regimes affect divergence, we generated additional simulations so that every simulation now has an identical paired population with the same starting conditions and parameters. We then asked how much total change these pairs have experienced and how much they have diverged.

Based on the adaptive responses we observed earlier (Fig 3) and on the harmfulness of environmental changes (Fig 4), we expected evolving with an antagonistic partner and highly directional abiotic change to result in a relatively high amount of evolution within a population and, as a consequence, high divergence between replicated populations.

We first looked at the how much the paired populations changed phenotypically (the scalar sum of phenotypic change over both populations, Fig 5A) and genetically (summed number of fixed mutations over both populations, Fig 5B). Directionality in abiotic populations behaved as expected; within each regime type, the more directional abiotic change was, the greater the genetic and phenotypic change. Also as expected, conflict of all three types caused more divergence than abiotic change of low directionality. Mutualism behaved in opposite fashion to conflict, with low divergence relative to its corresponding abiotic simulations.

**Figure 5:**
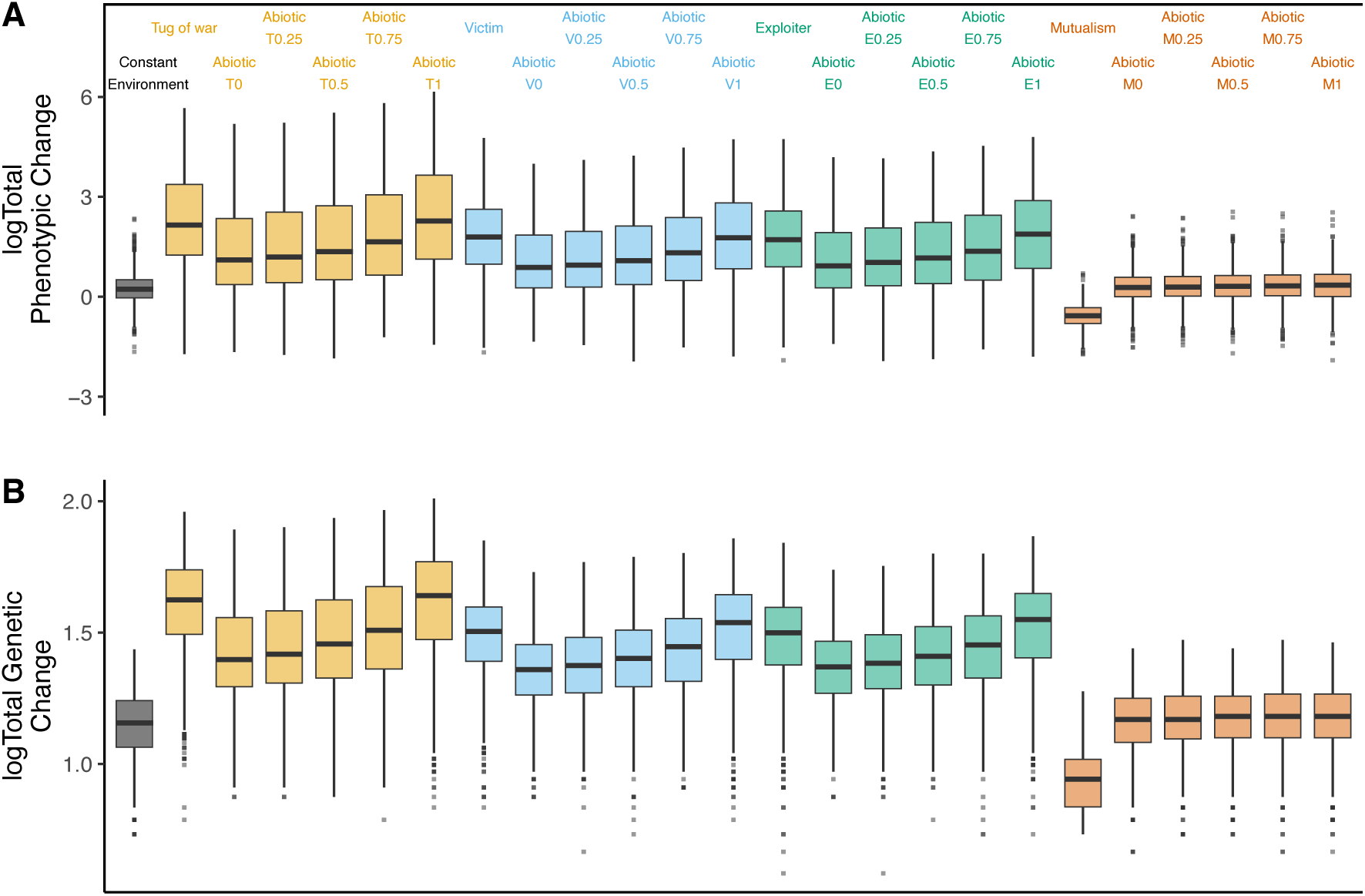
Conflict and directional change result in greater evolution. (A) Log transformed total phenotypic change for all environmental change regimes. Total phenotypic change is the summed scalar distance of all phenotypic changes over the course of a simulation across paired populations. (B) Log transformed total genetic change for all environmental change regimes. This is the total number of fixed mutations between population pairs. Colors and boxplots are defined as in Fig. 4.

The high genetic and phenotypic change seen in biotic and highly directional simulations might be expected to lead to high phenotypic divergence (calculated as the phenotypic distance separating paired populations at the end of a simulation). This was the case in abiotic simulations (Fig 6A). It was also the case for victims and exploiters. But surprisingly, this pattern was broken under tug-of-war conflict (Fig 6A). This pattern of high phenotypic change but relatively low phenotypic divergence makes sense because tug-of-war conflict involves frequent fixations but with evolution remaining restricted around the two parties’ fixed optimum values [3].

**Figure 6:**
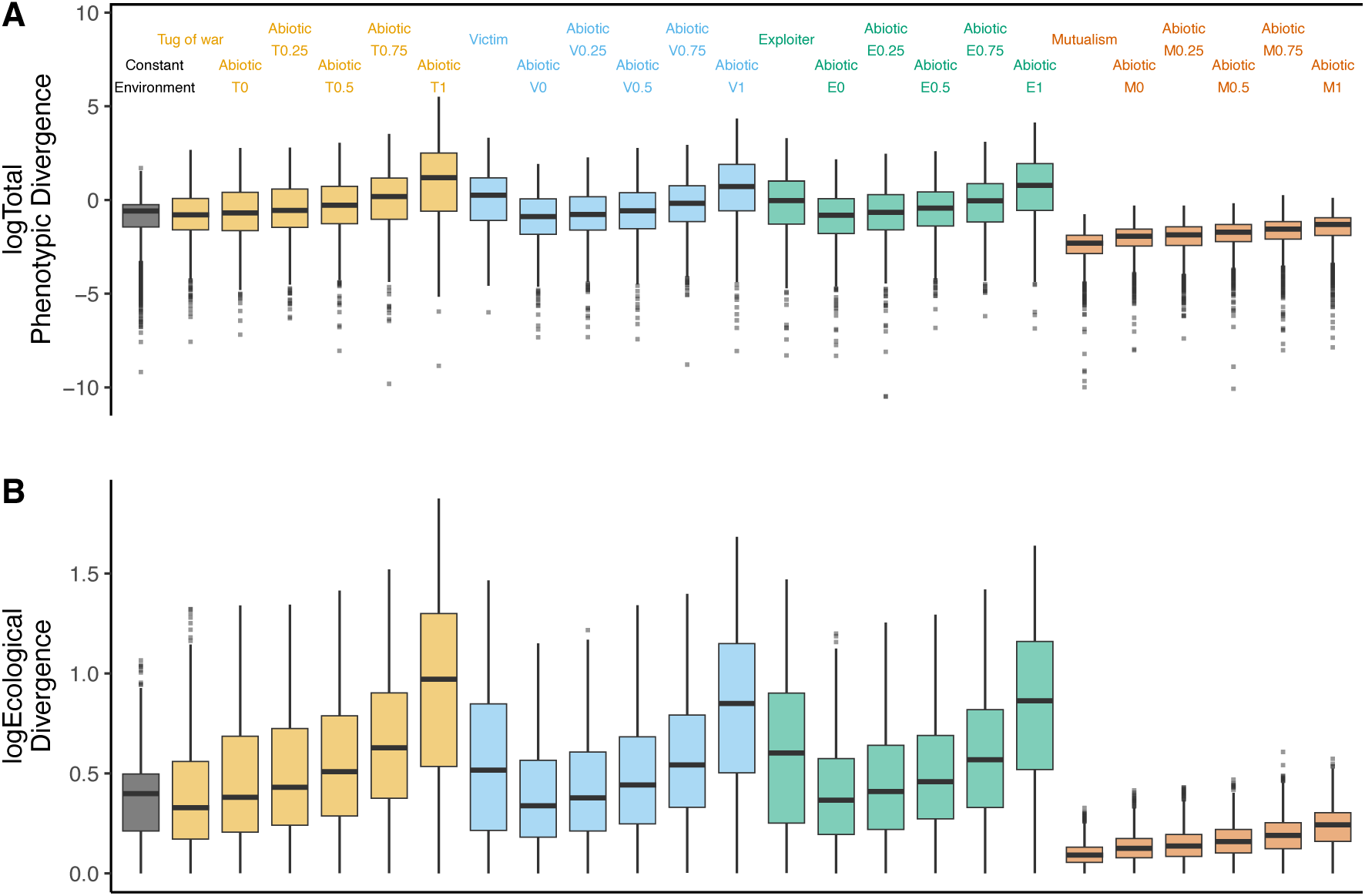
Conflict and directional change show greater phenotypic and ecological divergence, except for tug-of-war conflict. (A) Log transformed phenotypic divergence between paired populations for all environmental change regimes. Phenotypic divergence is the phenotypic distance between the populations at the end of the simulation. (B) Log transformed ecological divergence for all environmental change regimes. Ecological divergence is the distance between optima (abiotic) or partner traits (biotic) at the end of simulations. Colors and boxplots are defined as in Fig 4.

This is supported by the fact that the degree of phenotypic divergence is well predicted by the degree of ecological divergence, the distance between the final environments of the two populations (Fig 6B). In particular, both values are low for tug of war (where the ecological divergence is the distance between partner values).

### Directional abiotic change reduces migrant and hybrid fitness, but biotic conflict does not

One way that environmental change can affect speciation is through migrants failing to establish in different environments [51]. We tested how adapting to different environmental change regimes affected migrant fitness by simulating paired evolving populations with the same starting values experiencing the same environmental change regimes (though different instantiations of that regime). The fitness of migrants in a foreign environment was assessed relative to the fitness of local residents (Fig 7A shows an example with constant environment simulations that have adapted to two different optima).

**Fig 7:**
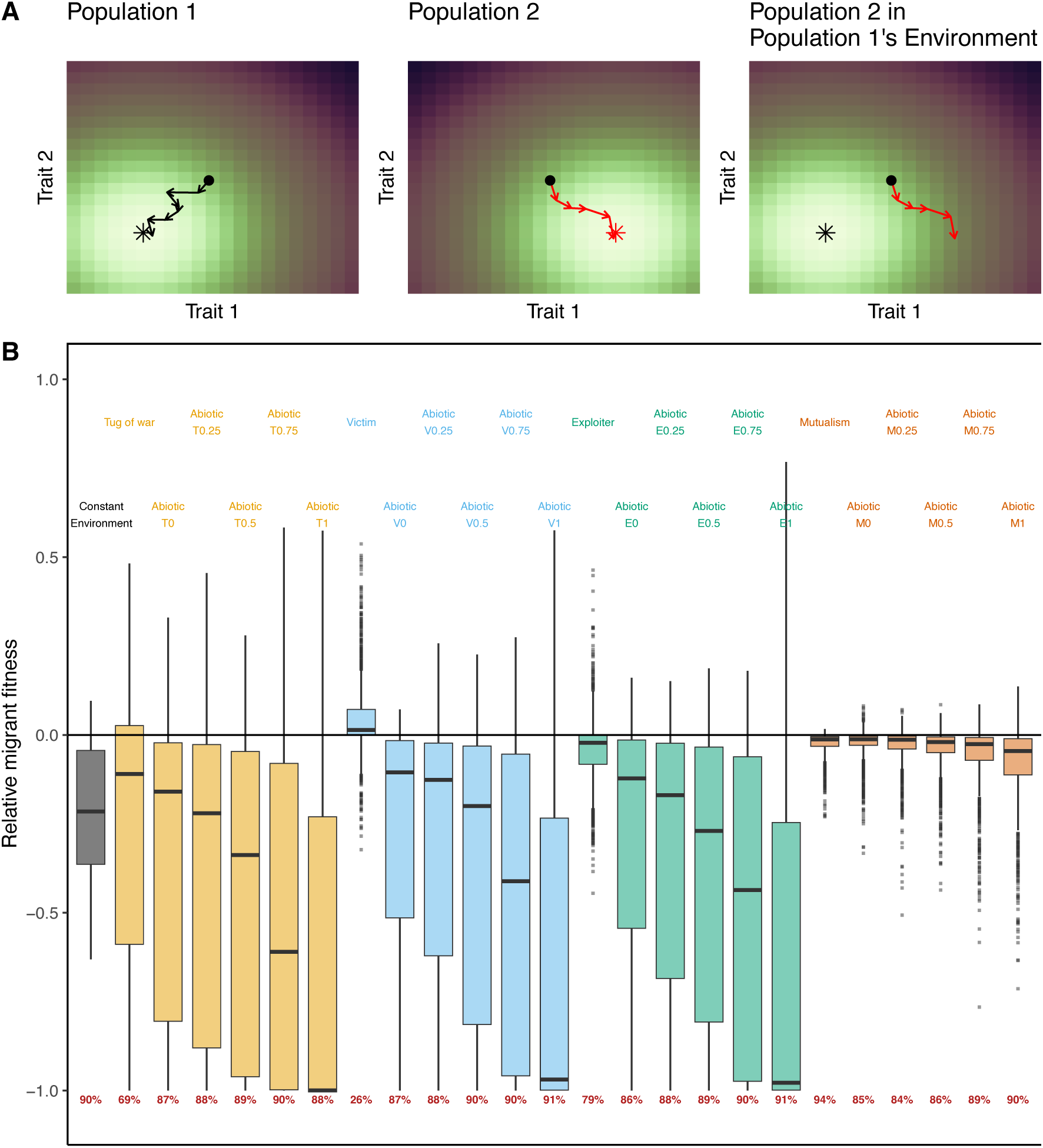
Migrants from environments with biotic conflict have a higher chance of success than migrants from other environmental change regimes. (A) Diagram of how migrant fitness was assessed using adaptation to two constant environments as an example. Black and red populations evolve separately and then the red phenotype is moved into the black environment. Relative migrant fitness shown in (B) is the selection coefficient for foreign (red in A) individuals relative to local (black in A) individuals, in the local black environment. Colors and boxplots are defined as in Fig 3. Red numbers at the bottom of B are the percent of migrants that are worse off than local residents (those with relative migrant fitness below the 0 line).

As expected, most migrants have reduced fitness relative to the local residents (Fig 7B). In the abiotic simulations, higher environmental directionality leads to lower migrant fitness, which is expected given that they have higher divergence. Mutualistic populations yielded relatively high migrant fitness, as expected from their low divergence.

However the patterns of fitness reduction were not always as expected under conflict. Strikingly, migrants in biotic conflict regimes tended to have higher relative fitness than migrants in their corresponding abiotic change simulations. This might be expected from the low divergence in the tug-of-war regime, but not from the high divergence of victims and exploiters. The most dramatic example of this comes from victim migrants that were worse off than local residents in only 26% of the simulations (red numbers at the bottom of Fig 7B).

Different environmental change regimes, and their concomitant changes to adaptive responses, should also affect the fitness of hybrids between diverging populations. We investigated this by generating hybrids between parents from different populations and measuring fitness of hybrid offspring [35,42]. We created F1 crosses using our paired local and foreign parents from above by adding together, at random, half of the fixed mutations in each parent (Fig 8A).

**Fig 8:**
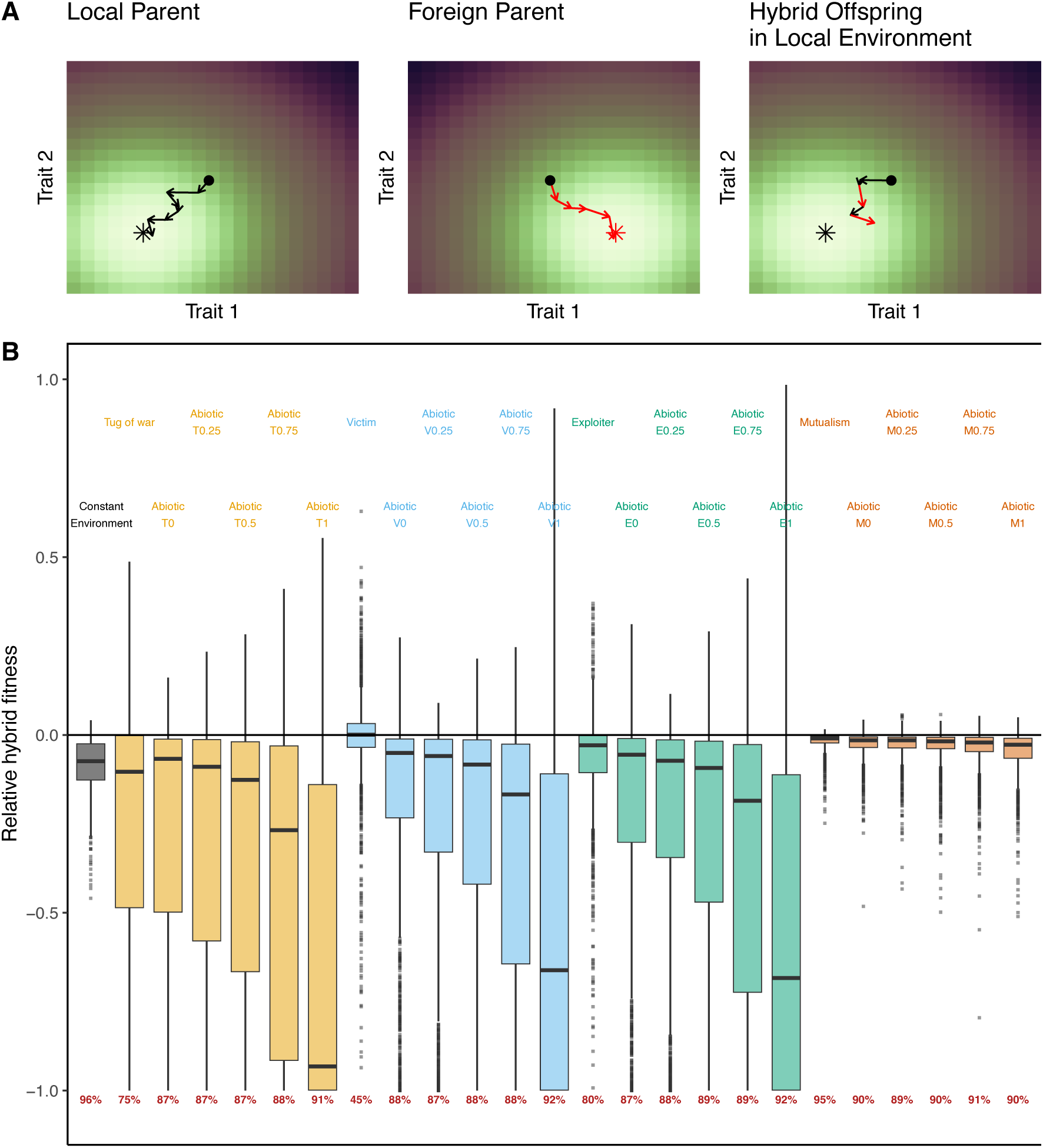
Hybrids from environments with biotic conflict have a higher chance of success than hybrids from other environmental change regimes. (A) Diagram of hybridization for populations adapting to different optima. Hybrids consist of 50:50 mixes of fixations from both parents. Relative hybrid fitness shown in (B) is the selection coefficient for hybrids (mixed black and red in A) populations relative to local (black in A) populations, in the local black environment. (B) Distributions of relative hybrid fitness of F1 crosses. Colors and boxplots are defined as in Fig 3. Red numbers at the bottom of B are the percent of hybrids that are worse off than their parents (those with relative hybrid fitness below the 0 line).

Hybrid fitness (Fig 8B) showed essentially the same patterns we described above for migrant fitness (Fig 7B). Most notably, hybrids in biotic simulations again were better off than their corresponding abiotic simulations (Fig 8B). The two measures of migrant and hybrid fitness are also positively correlated across all environmental change regimes, except in victims and exploiters, which had no strong correlation (Fig 9). Thus, conflict regimes were less likely to result in low migrant and hybrid fitness leading to speciation, despite their high rates of change via arms races.

**Fig 9:**
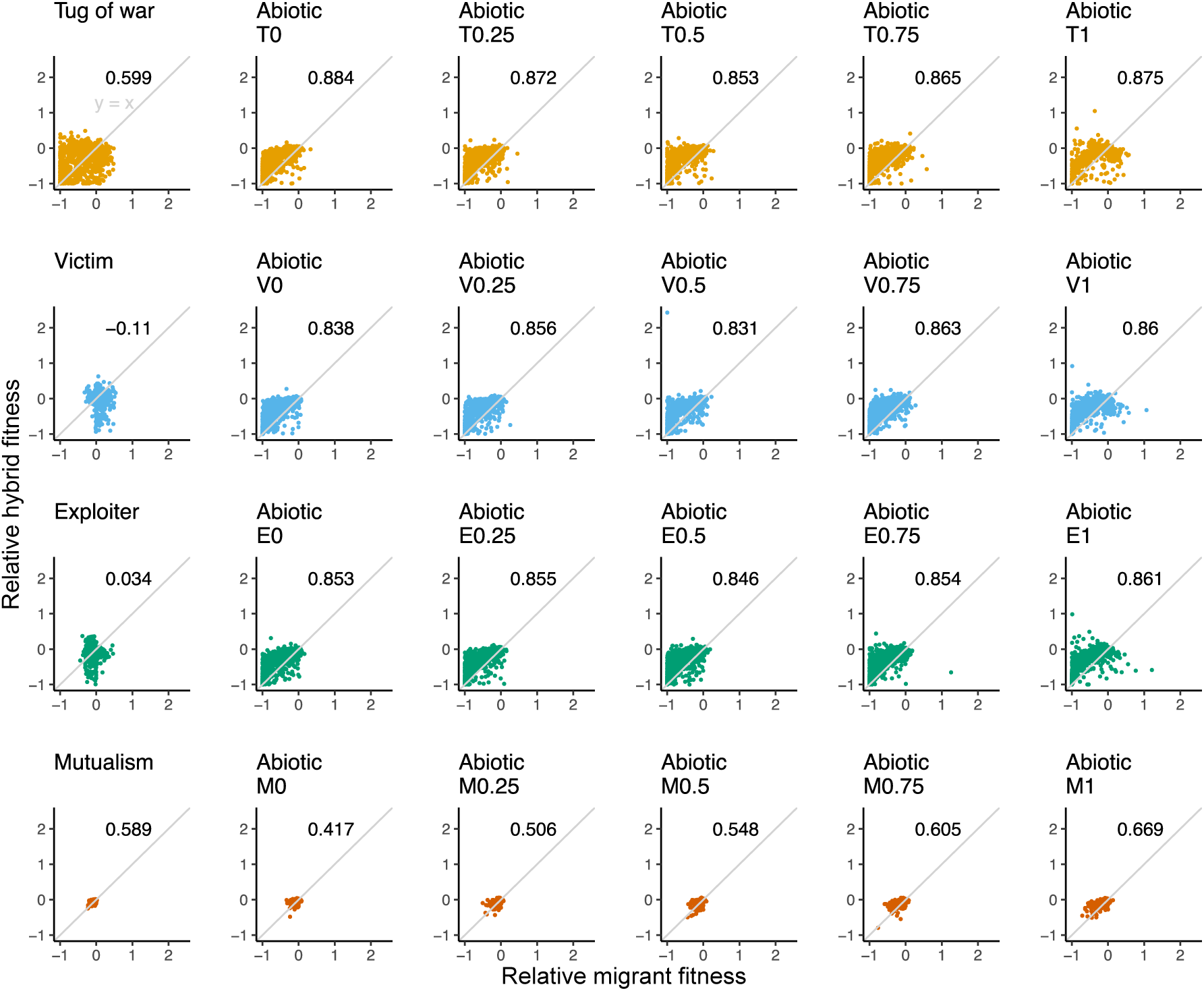
Hybrid and migrant fitness values are correlated except in victims and exploiters. Each panel shows individual relationships between relative migrant fitness and relative hybrid fitness. Pearson correlations are shown in the top right corner of each plot. Gray line is the identity line.

## Discussion

Adaptation and speciation are central questions in the study of evolution, but our knowledge of how these are affected by different environments is limited.

Adapting to a biotic environment is thought to be different than adapting to an abiotic environment [1–7], but the details of this idea have not been fully explored. Our previous modeling has identified some differences among the adaptive responses from tug-of-war conflict, uncorrelated abiotic change, and standard adaptation [3]. We extended these models to include multiple traits and added additional breadth with mutualisms, victim-exploiter conflict, and correlated abiotic change. For each environmental change regime, we also examined how much populations diverged and how this divergence affects migrant and hybrid fitness.

Our results show that different kinds of environmental change can lead to distinct adaptive responses and can affect fitness differences that cause speciation.

Surprisingly, we found that conflict may not be a great driver of speciation relative to changing abiotic environments.

### Different environmental regimes cause different adaptive responses

We found that, in terms of their adaptive responses, different ecological change regimes occupied informative positions in principal component space that corresponded with average fitness (Fig 3). In particular, conflict and strongly directional abiotic changes resulted in severe maladaptation on average while less directional abiotic change regimes resulted in more moderate maladaptation.

Mutualisms and adapting to a constant environment resulted in high amounts of adaptation. These patterns in adaptation and maladaptation affected downstream components of the adaptive response (Fig 3). As is well known, when a population has low fitness, it will fix more and larger fixations with large selection coefficients in order to adapt [3,8,49] and this pattern was recapitulated in our simulations (Fig 3). Our work now also demonstrates that these large and frequent fixations will tend to knock traits that are only being indirectly selected off of their optima (Fig 3) because of pleiotropy, as was found in another recent study [63].

We previously suggested that the differences between conflict and abiotic change were due to conflict resulting in changes that are consistently antagonistic [3]. We explicitly tested this by examining the distribution of fitness effects of the environment (DFEE) and found that conflict often results in negatively shifted DFEEs relative to low-directional abiotic change of the same magnitude (Fig 4).

### Conflict is a weak cause of speciation

We expected different kinds of environmental change to affect speciation potential by changing a population’s adaptive response and the amount of ecological and genetic divergence between populations. We expected that the high amount of change under conflict and highly directional abiotic selection would drive more population divergence and that greater divergence would drive more speciation (Fig 1) [6,21,29,52,53].

Abiotic evolution behaved as expected within each of the regimes where the amount of change was held constant (T, V, E, M). Higher directionality (correlation between environmental changes) cause more deleterious fitness changes (Fig 4) and larger total amounts of genetic and phenotypic changes (Fig 5). These larger changes resulted in more divergence between paired populations (Fig 6) and lower migrant and hybrid fitness (Figs 7&8). Correlated environmental change is therefore associated with greater speciation potential than uncorrelated change.

We reasoned that mutualism, where trait convergence is favored, would show the opposite pattern to that expected under an arms race with environmental changes being largely positive. he fraction of deleterious environmental changes was quite low and the average change was positive (Fig 4). Mutualism had less evolutionary change (Fig 5) and less population divergence at the end of the simulation than any other treatment (Fig 6). As expected from this, both migrant and hybrid fitness (Figs 7&8) were relatively high, though it should be noted that migrants and hybrids were almost always slightly worse off than their parents (94% of migrants and 95% of hybrids; Fig 7&8). This makes sense if parents are highly adapted to their local partners and migrants and hybrids end up with some amount of trait mismatch. Because divergence is low in mutualisms, the consequences of mismatch tend to have only small effects on migrant and hybrid fitness.

We found deviations from our hypotheses (Fig 1) when looking at the three types of biotic conflict evolution – tug of war, victim, and exploiter. They all showed the initial expected effects of conflict. Change in these biotic environments was highly detrimental to fitness (Fig 4) because their opponents are selected to improve their own fitness at the expense of their partners [3,6]. Their abiotic counterparts, where the amount of environmental change was the same, did not experience such deleterious effects except when the environmental changes were highly directional (Fig 4). Thus, conflict and highly directional change were “malicious;” they nearly always lowered fitness.

Populations with conflict also showed high rates of genetic and phenotypic change or arms races (Fig 5) as each partner tried to outrun the other, supporting hypothesis 1 in Fig 1. But these arms races did not lead to higher speciation propensity compared to abiotic populations undergoing the same amount of environmental change. Instead, they resulted in high migrant and hybrid fitness (Figs 7&8). The reasons for this are different for tug-of-war versus victim-exploiter conflict.

In victim-exploiter conflict, the hypothesis fails at step 3 of Fig 1; high divergence does not lead to low migrant and hybrid fitness. The primary reason is that victim and exploiter fitnesses are inversely related; a larger distance benefits victims and a smaller one benefits exploiters. The net effect of migration on distance is zero. To see this, let the distance between the victim in population *i* and the exploiter in population *j* be *D*(*V_i_,E_j_*). Now consider the four possible migrants between populations 1 and 2. First, if a victim in population 1 migrates to population 2, its distance from exploiters changes by *D*(*V_1_,E_2_*)-*D*(*V_1_,E_1_*). For a victim migrating from population 2, it is *D*(*V_2_,E_1_*)-*D*(*V_2_,E_2_*). For exploiters, the change in distance is either *D*(*V_2_,E_1_*)-*D*(*V_1_,E_1_*) or *D*(*V_1_,E_2_*)-*D*(*V_2_,E_2_*). The average distance change of the two victim migrators is exactly the same as the average change of the two exploiter migrators. Because a given change in distance benefits one class at the expense of the other, on average fitness gains and losses are equally likely.

This average does not necessarily mean that both victims and exploiters will have equal fitness gains and losses. If the two populations have moved far apart in trait space, moving to a new population will usually result in a greater distance, which would benefit a victim migrant and harm an exploiter migrant. Consistent with this, victim migrants lose fitness 26% of the time and exploiters 79% of the time (Fig 7). Given the logic above, we might expect on average losses between victims and exploiters to be around 50% since victims and exploiters will tend to alternate losses. This average for our data is close at 52.5%, but this number is slightly higher than the expected 50%. However, the argument above concerns only conflict traits, and our data sometimes include pleiotropic traits. If we look at the subset of simulations with no pleiotropic traits, we find an average value of 50.5%, much closer to the expected value if victims and exploiters trade gains and losses.

Tug-of-war conflict is different, though with the same end result of migrant and hybrid fitness being higher than expected. Here the hypothesis chain breaks down at hypothesis 2 of Fig 1, Although environmental changes are quite negative (Fig 4) and evolutionary responses are high changes (Fig 5), this does not translate into high divergence (Fig 6). This is because, in contrast to abiotic evolution where the optimum keeps changing, tug-of-war conflict occurs primarily in a fixed space between the optima of the two parties (though perhaps occasionally overshooting). There is a lot of evolution in that space because of conflict, but little divergence because in both populations there is constant pulling and pushing around a central value (the exact center in our case where the two parties have the same parameters). The tug of war causes evolutionary churn, which does not strongly promote speciation.

### Caveats and conclusions

Our focus here was on the different rates of diversification and speciation induced by different selective regimes, especially with respect to antagonistic interactions. Biotic and abiotic interactions can differ for various other reasons [52], not all of which we include. The escape-and-radiate hypothesis posits that diversification occurs not via antagonism itself, but when a species escapes its antagonist, for example an insect moving to a new food plant [25]. The geographic mosaic theory of coevolution contains arms-race ideas but is thought to also apply to mutualism [64]. Later models have suggested that mutualism is more likely to inhibit local adaptation and speciation [59,65]. We also do not include between-species competition, an antagonistic biotic regime that can promote ecological speciation via a different mechanism, character displacement [17]. Finally, we do not consider how biotic interactions may influence speciation by affecting gene flow, for example when pollinators disperse plant genes.

By setting aside these effects we focused on the question of how conflict and arms races affected speciation potential. Our models apply most directly to predator-prey and host-pathogen antagonisms. But the two coevolving gene sets in our models can also be viewed as two distinct sets within a species. They can therefore also apply to male-female conflict [66,67] or conflict due to selfish genetic elements [53,68].

It should also be noted that a genome consists of many genes that are all evolving according to different environmental change regimes. Genes involved in conflicts may represent only a small fraction of the genome. Speciation potential for a population will then involve the proportional consequences of these conflict genes along with genes evolving according to other pressures. Mutualisms may also involve some conflict and our mutualism models do not include this.

It is also important to remember that we have compared biotic and abiotic regimes that experience the same amount of environmental change. There is no particular expectation that this will be true in nature. If, for example, biotic environments change more rapidly in nature, that might rescue the idea that they could lead to more speciation. But it should be noted that the fixations of partner genes that cause biotic environmental change occur on the scale of many generations, whereas abiotic changes might occur on much shorter time scales. Ultimately this is an empirical question, but our models do show that biotic arms races include some factors that are unfavorable for speciation.

## Methods

### Fisher’s Geometric Model

To simulate Fisher’s geometric model, we created an R package called *SimGM* that is available from GitLab (https://gitlab.com/treyjscott/SimGM). Before describing the models with changing environments, we explain the standard geometric model where a haploid population adapts to a constant environment (Fig 2A). The geometric model assumes that a population with *n* total traits experiences stabilizing selection around some optimum phenotype **o** (we denote vectors in lowercase bold letters and matrices in uppercase bold letters). We assumed that fitness is a result of Gaussian stabilizing selection and is a function of the Euclidean distance *d* between the current phenotype **z** and the optimum **o**,

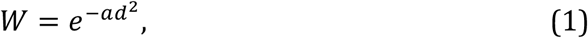

where *a* scales the strength of selection by making the fitness function narrower or wider for stronger and weaker selection, respectively. We assumed that the optimum is the origin for the purposes of explanation. We started simulations with phenotypes some distance from the origin such that populations are initially maladapted from the maximum possible fitness of 1 by an amount *L*, W _t = 0_ = 1 – *L*, where *L* is the lag load [69]. The lag load *L* was used in all models (though varied across replicates; see below) to control initial fitness [3]. Using equation 1 and W = 1-*L*, we solved for the Euclidean distance for a given lag load, 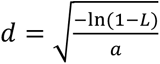.

Of our total of *n* traits, we now need to distinguish two classes. *v* of them are started away from their optimum and are selectively responsive to changes in the environment. The remaining *p = n* - *v* traits are started at their optimum and are selectively unresponsive (they have fixed optima). Once the initial Euclidean distance *d* is calculated, we ensure that this total distance is distributed among the v traits by making the initial value for each of the *v* traits equal to 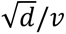.

We make the distinction between environmentally responsive and unresponsive traits to incorporate traits that are only pleiotropically affected by environmental changes. These *p* traits can be moved away from their optimum by selection on other traits, and they can be important for speciation [35]. We use this same method of having *p* unresponsive traits start at their optimum (the origin) for all models to easily compute the amount unresponsive traits are knocked off their optimum by selection on responsive traits. The amount of deleterious pleiotropic change is calculated as the Euclidean distance of these pleiotropic traits from the origin at the end of simulations.

In order for a population to adapt and move towards the optimum, it must fix beneficial mutations that change and improve the population’s current phenotype. Mutations are vectors in trait space that are added to the current phenotype and causing fitness to change to *w_mut_*. Mutations fix according to their selection coefficients *s = w_mut_/w_anc_ – 1*, where *w_anc_* is the fitness of the pre-mutation ancestor. For finite haploid populations with *N* individuals, a mutation fixes with probability

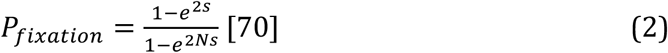

When a mutation is not fixed, the current phenotype remains unchanged. Since we are focused on the effects of selection in different changing environments, we set *N* to 10,000 to minimize the effects of drift. We assume that the strength of selection is strong and the mutation rate is low to ensure that adaptation can be modeled as a process of repeatedly drawing a single mutation, testing whether it fixes, and adding the fixed mutation to the current phenotype [71].

To control the size of mutations, we draw mutation total effect sizes *r* from an exponential distribution with mean 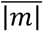 To convert this magnitude *r* into a random vector with *n* elements, we randomly drew angular coordinates and translated them from a spherical coordinate system to a Euclidean coordinate system [48,72].

### Abiotic Change

To incorporate changing abiotic environments, the optimum position is shifted over time [3,8,38–41]. Changing abiotic environments can be thought of as a continuum, where at one end there is change that is uncorrelated (directionally random; Fig 2B) and at the other end change is completely autocorrelated (the direction of future changes is the same as the past; Fig 2C). Most natural environments likely occupy some intermediate position on this continuum.

To compare to simulations with biotic change, we make the size and frequency of abiotic changes equal to those from the *v* environmentally responsive traits from a comparison biotic change simulation with the same parameter values (see Comparing environmental change regimes). Only the optima for the *v* environmentally responsive traits change during abiotic change simulations. To control the size of changes to the optimum position, we use spherical coordinates to translate a magnitude to a vector to add to the optimum. We controlled the direction of change (which emerges naturally in biotic simulations) by defining the autocorrelation parameter *r*, the probability that a change at time *t* occurs in the same direction as the change at time *t-1.* Directions in this case are determined by the angles used to translate spherical coordinates to Euclidean coordinates [48,72]. If the direction changes, new angles are drawn randomly. We use draws from a Bernoulli distribution with *p* = *r* to determine whether changes occur in the same (1) or different (0) direction. We note that when *r* is 0, we obtain uncorrelated (random) change.

### Tug-of-war conflict

Biotic conflict between two parties has been modeled using a tug of war (Fig 2D) in the geometric model using joint phenotypes [3]. Joint phenotypes are shared by two interacting parties, where evolution in both sides affects the single joint phenotype [13]. Most interactions can be modeled as a joint phenotype. For example, in predatory garter snakes that feed on toxic newts [73], the probability that a snake can successfully eat a newt is a joint phenotype. It is a single measurable phenotype that can change due to evolution in the garter snake population or in the newt population. Evolution with a single tug of war in the geometric model results in long-term maladaptation due to Sisyphean or tug-of-war arms races, where both sides are kept from their optimum values [3].

We can include evolutionary conflict over a joint phenotype with a tug-of-war model like we did for adaptation to a constant optimum, with the same fitness and fixations functions and same mutation distributions. However, now with two parties that are identical in every respect except for having different optima for their joint phenotype. We assumed that *v* of the *n* traits are joint phenotype traits involved in conflict between two parties. These *v* traits are the ones that are responsive to change in the environment, which now takes the form of evolutionary change in the other party’s genes. The remaining n - *v* pleiotropic traits, are initially at their optimum and are only indirectly affected by conflict.

The *v* mutations in both parties affect the tug of war over the joint phenotype, but parties have different optimal values where fitness is maximized [3]. Because we made both parties the same except for the position of their optima, conflict in these simulations is symmetrical. For convenience, we started joint phenotype traits at the origin and with the parties’ optima equal distances away on opposite sides. Thus, movement from the origin towards one optimum normally moves the other party farther from its optimum. The initial distance to the optimum, and individual optimum trait values, is calculated as a function of initial lag load as was done for starting phenotypes in the standard adaptation model. These changes are for convenience as our results are invariant to the actual locations in trait space.

To accommodate two parties jointly affecting a phenotype, for each time step (iteration), we randomly choose one party to draw and potentially fix a mutation first before the other party then has a turn in that same time step. This ensures that both sides have equal opportunities to affect the joint phenotype.

### Victim-Exploiter conflict

Tug-of-war conflict results in escalatory arms races where both sides fix large back-and-forth mutations [3]. Biotic conflicts could alternatively involve trait matching in which one party tries to evolutionarily evade the chase of the other party. Victim-exploiter conflict (Fig 2E) is a well-studied example of trait-matching in quantitative genetics [15,16,60], where exploiters get a fitness benefit from matching their traits to those of victims while victims experience reduced fitness when traits match. For example, brood parasites attempt to match their egg and chick phenotypes to those of their hosts in order to secure food at the expense of the host’s own offspring [74]. Another less literal example of matching is when pathogens are able to exploit hosts by binding to host proteins and causing infections [75]. Over evolutionary time, matching results in exploiters chasing victims in trait space.

The basic algorithm for victim-exploiter conflict is like that of tug-of-war conflict in that both parties take turns drawing and potentially fixing mutations.

However, in contrast to a single tug of war, victims and exploiters have their own conflict traits that interact without stabilizing selection.

Exploiters gain a fitness benefit when their traits match those of the victim, while victims have lower fitness [15,60]. The probability that exploiters match victims is

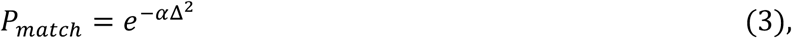

where α determines how important trait values are for determining the outcome of the interaction and Δ is the distance between *v* victim and exploiter traits that are involved with conflict [16]. We set α to 6, an intermediate value that ensures populations will coevolve. We also note that just like in tug-of-war conflict simulations, the conflict traits used to calculate Δ can be a subset defined by *v* (the number of traits that are selectively responsive to the conflict). The remaining *n* – *v* = *p* traits are not involved in conflict but can be selected when pushed off their optima by pleiotropy.

Fitness for victims and exploiters is then the product of fitness due to stabilizing selection on unresponsive traits (W_stab_) and fitness due to the interaction [16]:

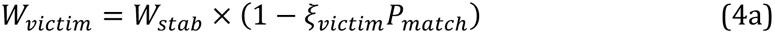

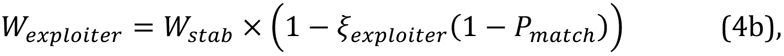

where *ξ_victim_* and *ξ_exploiter_* are the fitness costs and benefits of the interaction. To ensure that exploiter fitness could not exceed 1, we included a negative *ξ_exploiter_* term. For simplicity, we ensure that both victims and exploiters experience the same fitness incentives by assuming that the costs and benefits of matching are the same for victims and exploiters such that *ξ_victim_* = *ξ_exploiter_*. These fitness equations can be used to calculate selection coefficients and determine whether new mutations fix.

### Mutualism

Mutualisms were modeled like victim-exploiter evolution with trait matching (Fig 2F), except that matching benefits both species instead of benefiting one and harming the other [16]. Both parties receive a benefit from the interaction:

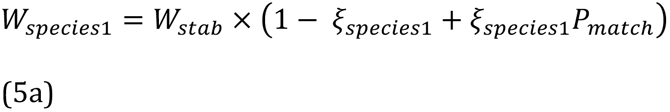

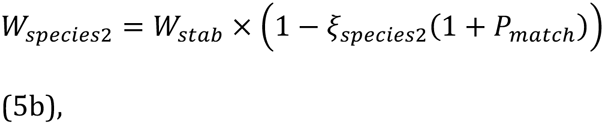

where *P_match_* is the trait matching equation for victims and exploiters. We again subtract the benefit to ensure that the maximum fitness is 1 and assume that both sides experience the same fitness consequences so that *ξ_species_*_1_ = *ξ_species_*_2_.

### Comparing environmental change regimes

In nature, biotic and abiotic changes occur with different frequencies and magnitudes. Populations also differ in how maladapted they are at the start of environmental change. However, we aim to identify signatures of adaptation and divergence for different kinds of environmental change by controlling for these differences as much as possible. We will first discuss making initial lag loads equal and then making the environmental changes equal.

For tug-of-war conflict, we can directly control the amount of initial lag load and make it equal to that of standard adaptation simulations (see *Tug-of-war conflict* above). For victim-exploiter and mutualisms, we start populations so that both sides match half of the time. Because *P_match_* depends on the distance between victim and exploiter traits, we can rearrange the *P_match_* equation (equation 3) in terms of the distance, 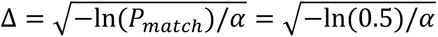, and then use this distance when creating starting trait values. For convenience, we start victim and exploiter conflict traits equidistant across from the origin (with the total distance between them being Δ).

To make abiotic environmental change equal to change from conflict simulations, we changed the environment in each abiotic simulation (the optimum position) by the same magnitude that the environment changed in a corresponding (same parameters) biotic simulation for only environmentally responsive traits. If a partner mutation fixed in an abiotic simulation, which corresponds to a change in environment, then the optimum of a matching abiotic simulation was shifted by that amount but with a direction that was either random (r=0) or partially correlated with its previous direction change (r>0). If no mutation was fixed at a specific time during a biotic conflict simulation, then the optimum is not shifted in abiotic change simulations. This ensures that both biotic and abiotic simulations experience the same magnitude and timing of environmental change but not the same direction [3]. Because the biotic models showed differences in fixation sizes, we ran separate corresponding abiotic simulations for each kind of conflict. Thus, each kind of conflict change (tug of war, victim, exploiter, mutualism) has a corresponding set of five abiotic change simulations, one for each degree of autocorrelation.

To control for the effect of ecological divergence when the environments are constant, we simulate two standard adaptation populations that start with the same phenotype but adapt to different optima that do not change over time. The optimum of one population is the origin. For the second population, we randomly generate an optimum using spherical coordinates (same as how mutations are generated as described in the Fisher’s Geometric Model section) that ensures that both populations have the same amount of initial maladaptation.

### Migrant fitness

To test how different kinds of environmental change impact migrant inviability, we used pairs of parental populations that were simulated with identical parameters. We assigned one parental population to represent the local population and one to represent the foreign population. We calculated the fitness of the foreign phenotype in the local environment (the local optimum for abiotic, the local coevolving partner for biotic). This is straightforward using the fitness equations above except in the case of conflict with a tug of war. In this case, the joint phenotype is affected by evolution in both parties and the joint phenotype value in the foreign environment is not the same as the joint phenotype in the local environment. To account for this, we construct the joint phenotype that would result from an association between the foreign migrant and its new partner, the local antagonist, by adding the fixations of those two parties together, and then assessing the fitness of the migrant using its fitness function.

### Hybrid Crosses

Using the population pairs that were used to access migrant fitness, we constructed hybrid crosses by randomly combining fixations from both parents into an offspring phenotype. Since parental populations are monomorphic, parents have the same phenotype as their population means. Because our populations are haploid, offspring receive half of each parents’ fixations drawn at random. These fixations are added to the starting phenotype of the parent populations (before they diverged) to construct the offspring F1 phenotypes [35,42].

Once offspring phenotypes have been constructed, fitness can be calculated in either parental environments according to the fitness functions. Conflict over joint phenotypes requires summing the fixations passed to hybrids with the fixations of antagonists in the test environment to correctly calculate the joint phenotype and hybrid fitness. This summing with the antagonist is necessary because the phenotype is jointly constructed by both parties.

### Measures from diverging parental populations

In addition to measures of the adaptive response of individual population simulations, we also calculated important outcome variables at the end of simulations from paired populations that had the same parameters and initial starting conditions. These measures captured how deleterious environmental changes were, how much phenotypic evolution occurred, how different environments were, and how genetically diverged populations were. We called these measures the fitness effect of environmental change, total genetic change, total phenotypic change, phenotypic divergence, and ecological divergence.

To understand how environmental change affected fitness, we calculated individual selection coefficients for biotic and abiotic environmental changes. To do this we modified the usual selection coefficient equation from above to be, *s_dfee_ = w_after_/w_before_ – 1*, where *w_before_* is the fitness of the population before the environmental change and *w_after_* is the fitness of the population after the environmental change. To summarize the distribution of fitness effects of the environment, we calculated the median fitness effect and the proportion of fitness effects that were harmful (*s_dfee_* < 0).

To measure total genetic change in our simulations, we summed the number of fixed mutations from both populations. To measure total phenotypic change, we summed the magnitude (scalar sum) of all fixed mutations from both populations in a simulation.

For phenotypic divergence, we measure the Euclidean distance between the phenotypes in the two population at the end of the simulation. Because many changes can happen to phenotypes without a large effect on the final phenotypes, the total genetic change and phenotypic divergence may be quite different. To measure ecological divergence in our simulations, we similarly calculated the Euclidean distances of the environments from paired populations at the end of simulations. For abiotic environments, the ecological divergence was the distance between optimum values at the end of a simulation. For biotic environments, the ecological divergence was the distance between partner traits from environments 1 and 2 because partners constitute the environments in the two populations. For example in victims, the ecological divergence would be the distance between exploiter traits (the victim’s environment) in both paired simulations.

### Statistical Analysis

To visualize differences in adaptive responses between models of environmental change, we performed principal components analysis using the *prcomp* function in R. All analyses and simulations were performed in R version 4.3.3 [76].

## Acknowledgements

We thank Israt Jahan for comments on an early draft of this manuscript and students in the Population Genetics class at Washington University in St. Louis for testing the SimGM program.

## Notes

### Competing Interest Statement

The authors have declared no competing interest.

### Funding

- NSF: DEB 2237266 & NSF DEB-1753743

## References

1. Agrawal AF, Whitlock MC. Environmental duress and epistasis: how does stress affect the strength of selection on new mutations? Trends Ecol Evol. 2010;25: 450–458. doi:10.1016/j.tree.2010.05.003

2. Louthan AM, Kay KM. Comparing the adaptive landscape across trait types: larger QTL effect size in traits under biotic selection. BMC Evol Biol. 2011;11: 60. doi:10.1186/1471-2148-11-60

3. Scott TJ, Queller DC. Long-term evolutionary conflict, Sisyphean arms races, and power in Fisher’s geometric model. Ecol Evol. 2019;9: 11243–11253. doi:10.1002/ece3.5625

4. Whiteman NK. Evolution in small steps and giant leaps. Evolution. 2022; evo.14432. doi:10.1111/evo.14432

5. Baskett CA, Schemske DW. Evolution and genetics of mutualism. Mutualism. Oxford University Press Oxford, UK; 2015. pp. 77–93.

6. Queller DC, Strassmann JE. Evolutionary Conflict. Annu Rev Ecol Evol Syst. 2018;49: 73–93. doi:10.1146/annurev-ecolsys-110617-062527

7. Scott TJ. The evolutionary potential of symbiosis. J Evol Biol. 2025; voaf152. doi:10.1093/jeb/voaf152

8. Matuszewski S, Hermisson J, Kopp M. Fisher’s geometric model with a moving optimum. Evolution. 2014;68: 2571–2588. doi:10.1111/evo.12465

9. O’Brien AM, Jack CN, Friesen ML, Frederickson ME. Whose trait is it anyways? Coevolution of joint phenotypes and genetic architecture in mutualisms. Proc R Soc B Biol Sci. 2021;288: 20202483. doi:10.1098/rspb.2020.2483

10. Coyne JA, Orr HA. Speciation. Sinauer associates Sunderland, MA; 2004.

11. Nosil P. Ecological speciation. Oxford University Press; 2012.

12. Dawkins R, Krebs JR. Arms races between and within species. Proc R Soc Lond B Biol Sci. 1979;205: 489–511.

13. Queller DC. Joint phenotypes, evolutionary conflict and the fundamental theorem of natural selection. Philos Trans R Soc B Biol Sci. 2014;369: 20130423. doi:10.1098/rstb.2013.0423

14. Vasseur DA, Yodzis P. The color of environmental noise. Ecology. 2004;85: 1146–1152. doi:10.1890/02-3122

15. Gavrilets S. Coevolutionary Chase in Exploiter–Victim Systems with Polygenic Characters. J Theor Biol. 1997;186: 527–534. doi:10.1006/jtbi.1997.0426

16. Nuismer S. Introduction to coevolutionary theory. Macmillan Higher Education; 2017.

17. Schluter D. The ecology of adaptive radiation. OUP Oxford; 2000.

18. Nosil P. Ecological speciation. Oxford University Press; 2012.

19. Polato NR, Gill BA, Shah AA, Gray MM, Casner KL, Barthelet A, et al. Narrow thermal tolerance and low dispersal drive higher speciation in tropical mountains. Proc Natl Acad Sci. 2018;115: 12471–12476. doi:10.1073/pnas.1809326115

20. Thompson JN. The coevolutionary process. University of Chicago press; 1994.

21. Presgraves DC. The molecular evolutionary basis of species formation. Nat Rev Genet. 2010;11: 175–180. doi:10.1038/nrg2718

22. Bérénos C, Schmid-Hempel P, Wegner KM. Antagonistic Coevolution Accelerates the Evolution of Reproductive Isolation in *Tribolium castaneum*. Am Nat. 2012;180: 520–528. doi:10.1086/667589

23. Hembry DH, Weber MG. Ecological Interactions and Macroevolution: A New Field with Old Roots. Annu Rev Ecol Evol Syst. 2020;51: 215–243. doi:10.1146/annurev-ecolsys-011720-121505

24. Coughlan JM. The role of conflict in shaping plant biodiversity. New Phytol. 2023; nph.19233. doi:10.1111/nph.19233

25. Ehrlich PR, Raven PH. Butterflies and plants: a study in coevolution. Evolution. 1964; 586–608.

26. Edger PP, Heidel-Fischer HM, Bekaert M, Rota J, Glöckner G, Platts AE, et al. The butterfly plant arms-race escalated by gene and genome duplications. Proc Natl Acad Sci. 2015;112: 8362–8366. doi:10.1073/pnas.1503926112

27. Allio R, Nabholz B, Wanke S, Chomicki G, Pérez-Escobar OA, Cotton AM, et al. Genome-wide macroevolutionary signatures of key innovations in butterflies colonizing new host plants. Nat Commun. 2021;12: 354. doi:10.1038/s41467-020-20507-3

28. Kawahara AY, Storer C, Carvalho APS, Plotkin DM, Condamine FL, Braga MP, et al. A global phylogeny of butterflies reveals their evolutionary history, ancestral hosts and biogeographic origins. Nat Ecol Evol. 2023 [cited 16 May 2023]. doi:10.1038/s41559-023-02041-9

29. Brucker RM, Bordenstein SR. Speciation by symbiosis. Trends Ecol Evol. 2012;27: 443–451. doi:10.1016/j.tree.2012.03.011

30. Xu S, Schlüter PM, Scopece G, Breitkopf H, Gross K, Cozzolino S, et al. Floral isolation is the main reproductive barrier among closely related sexually deceptive orchids. Evolution. 2011;65: 2606–2620. doi:10.1111/j.1558-5646.2011.01323.x

31. Wessinger CA, Katzer AM, Hime PM, Rausher MD, Kelly JK, Hileman LC. A few essential genetic loci distinguish Penstemon species with flowers adapted to pollination by bees or hummingbirds. Barton NH, editor. PLOS Biol. 2023;21: e3002294. doi:10.1371/journal.pbio.3002294

32. Brady SP, Bolnick DI, Barrett RDH, Chapman L, Crispo E, Derry AM, et al. Understanding Maladaptation by Uniting Ecological and Evolutionary Perspectives. Am Nat. 2019;194: 495–515. doi:10.1086/705020

33. Seehausen O, Butlin RK, Keller I, Wagner CE, Boughman JW, Hohenlohe PA, et al. Genomics and the origin of species. Nat Rev Genet. 2014;15: 176–192. doi:10.1038/nrg3644

34. Bradshaw HD, Schemske DW. Allele substitution at a flower colour locus produces a pollinator shift in monkeyflowers. Nature. 2003;426: 176–178. doi:10.1038/nature02106

35. Yamaguchi R, Otto SP. Insights from Fisher’s geometric model on the likelihood of speciation under different histories of environmental change. Evolution. 2020;74: 1603–1619. doi:10.1111/evo.14032

36. Thompson KA. Experimental Hybridization Studies Suggest That Pleiotropic Alleles Commonly Underlie Adaptive Divergence between Natural Populations. Am Nat. 2020;196: E16–E22. doi:10.1086/708722

37. Fisher RA. The genetical theory of natural selection. Clarendon Press; 1930.

38. Kopp M, Hermisson J. The Genetic Basis of Phenotypic Adaptation I: Fixation of Beneficial Mutations in the Moving Optimum Model. Genetics. 2009;182: 233–249. doi:10.1534/genetics.108.099820

39. Kopp M, Hermisson J. The Genetic Basis of Phenotypic Adaptation II: The Distribution of Adaptive Substitutions in the Moving Optimum Model. Genetics. 2009;183: 1453–1476. doi:10.1534/genetics.109.106195

40. Gordo I, Campos PRA. Evolution of clonal populations approaching a fitness peak. Biol Lett. 2013;9: 20120239. doi:10.1098/rsbl.2012.0239

41. Connallon T, Clark AG. The distribution of fitness effects in an uncertain world. Evolution. 2015;69: 1610–1618. doi:10.1111/evo.12673

42. Barton NH. The role of hybridization in evolution. Mol Ecol. 2001;10: 551–568. doi:10.1046/j.1365-294x.2001.01216.x

43. Fraïsse C, Gunnarsson PA, Roze D, Bierne N, Welch JJ. The genetics of speciation: Insights from Fisher’s geometric model: FISHER’S GEOMETRIC MODEL AND SPECIATION. Evolution. 2016;70: 1450–1464. doi:10.1111/evo.12968

44. Simon A, Bierne N, Welch JJ. Coadapted genomes and selection on hybrids: Fisher’s geometric model explains a variety of empirical patterns. Evol Lett. 2018;2: 472–498. doi:10.1002/evl3.66

45. Yamaguchi R, Wiley B, Otto SP. The phoenix hypothesis of speciation. Proc R Soc B. 2022;289: 20221186.

46. Kulmuni J, Wiley B, Otto SP. On the fast track: hybrids adapt more rapidly than parental populations in a novel environment. Evol Lett. 2023; qrad002. doi:10.1093/evlett/qrad002

47. Schneemann H, De Sanctis B, Welch JJ. Fisher’s Geometric Model as a Tool to Study Speciation. Cold Spring Harb Perspect Biol. 2024;16: a041442. doi:10.1101/cshperspect.a041442

48. Orr HA. The population genetics of adaptation: the distribution of factors fixed during adaptive evolution. Evolution. 1998;52: 935–949. doi:10.1111/j.1558-5646.1998.tb01823.x

49. Tenaillon O. The Utility of Fisher’s Geometric Model in Evolutionary Genetics. Annu Rev Ecol Evol Syst. 2014;45: 179–201. doi:10.1146/annurev-ecolsys-120213-091846

50. Huber CD, Kim BY, Marsden CD, Lohmueller KE. Determining the factors driving selective effects of new nonsynonymous mutations. Proc Natl Acad Sci. 2017;114: 4465–4470. doi:10.1073/pnas.1619508114

51. Nosil P, Vines TH, Funk DJ. Reproductive isolation caused by natural selection against immigrants from divergent habitats. Evolution. 2005;59: 705–719. doi:10.1111/j.0014-3820.2005.tb01747.x

52. Althoff DM, Segraves KA, Johnson MTJ. Testing for coevolutionary diversification: linking pattern with process. Trends Ecol Evol. 2014;29: 82–89. doi:10.1016/j.tree.2013.11.003

53. Crespi B, Nosil P. Conflictual speciation: species formation via genomic conflict. Trends Ecol Evol. 2013;28: 48–57. doi:10.1016/j.tree.2012.08.015

54. Paterson S, Vogwill T, Buckling A, Benmayor R, Spiers AJ, Thomson NR, et al. Antagonistic coevolution accelerates molecular evolution. Nature. 2010;464: 275–278. doi:10.1038/nature08798

55. Schulte RD, Makus C, Hasert B, Michiels NK, Schulenburg H. Multiple reciprocal adaptations and rapid genetic change upon experimental coevolution of an animal host and its microbial parasite. Proc Natl Acad Sci. 2010;107: 7359– 7364. doi:10.1073/pnas.1003113107

56. Obbard DJ, Welch JJ, Kim K-W, Jiggins FM. Quantifying Adaptive Evolution in the Drosophila Immune System. Begun DJ, editor. PLoS Genet. 2009;5: e1000698. doi:10.1371/journal.pgen.1000698

57. Enard D, Cai L, Gwennap C, Petrov DA. Viruses are a dominant driver of protein adaptation in mammals. eLife. 2016. doi:10.7554/eLife.12469

58. Buckling A, Rainey PB. The role of parasites in sympatric and allopatric host diversification. Nature. 2002;420: 496–499. doi:10.1038/nature01164

59. Yoder JB, Nuismer SL. When Does Coevolution Promote Diversification? Am Nat. 2010;176: 802–817. doi:10.1086/657048

60. Gilman RT, Nuismer SL, Jhwueng D-C. Coevolution in multidimensional trait space favours escape from parasites and pathogens. Nature. 2012;483: 328–330. doi:10.1038/nature10853

61. Lourenço J, Galtier N, Glémin S. Complexity, pleiotropy, and the fitness effect of mutations. Evolution. 2011;65: 1559–1571. doi:10.1111/j.1558-5646.2011.01237.x

62. Martin G, Lenormand T. A general multivariate extension of Fisher’s geometric model and the distribution of mutation fitness effects across species. Evolution. 2006;60: 15.

63. Rautiala P, Gardner A. The geometry of evolutionary conflict. Proc R Soc B Biol Sci. 2023;290: 20222423. doi:10.1098/rspb.2022.2423

64. Thompson JN. The geographic mosaic of coevolution. University of Chicago Press; 2005.

65. Kiester AR, Lande R, Schemske DW. Models of Coevolution and Speciation in Plants and Their Pollinators. Am Nat. 1984;124: 220–243. doi:10.1086/284265

66. Gavrilets S. Rapid evolution of reproductive barriers driven by sexual conflict. Nature. 2000;403: 886–889.

67. Parker G, Partridge L. Sexual conflict and speciation. Philos Trans R Soc Lond B Biol Sci. 1998;353: 261–274.

68. Brown JD, O’Neill RJ. Chromosomes, conflict, and epigenetics: chromosomal speciation revisited. Annu Rev Genomics Hum Genet. 2010;11: 291–316.

69. Maynard Smith J. What Determines the Rate of Evolution? Am Nat. 1976;110: 331–338. doi:10.1086/283071

70. Kimura M. The neutral theory of molecular evolution. Cambridge University Press; 1983.

71. McCandlish DM, Stoltzfus A. Modeling Evolution Using the Probability of Fixation: History and Implications. Q Rev Biol. 2014;89: 225–252. doi:10.1086/677571

72. Blumenson LE. A Derivation of n-Dimensional Spherical Coordinates. Am Math Mon. 1960;67: 63. doi:10.2307/2308932

73. Brodie ED, Brodie ED. Tetrodotoxin resistance in garter snakes: an evolutionary response of predators to dangerous prey. Evolution. 1990;44: 651–659. doi:10.1111/j.1558-5646.1990.tb05945.x

74. Feeney WE, Welbergen JA, Langmore NE. Advances in the Study of Coevolution Between Avian Brood Parasites and Their Hosts. Annu Rev Ecol Evol Syst. 2014;45: 227–246. doi:10.1146/annurev-ecolsys-120213-091603

75. Daugherty MD, Malik HS. Rules of Engagement: Molecular Insights from Host- Virus Arms Races. Annu Rev Genet. 2012;46: 677–700. doi:10.1146/annurev-genet-110711-155522

76. R Core Team. R: A language and environment for statistical computing. 2013.

